# Inefficient exploitation of accessory receptors reduces the sensitivity of chimeric antigen receptors

**DOI:** 10.1101/2021.10.26.465853

**Authors:** Jake Burton, Jesús A. Siller-Farfán, Johannes Pettmann, Benjamin Salzer, Mikhail Kutuzov, P. Anton van der Merwe, Omer Dushek

**Affiliations:** Sir William Dunn School of Pathology, University of Oxford, OX1 3RE, Oxford, UK

## Abstract

Chimeric antigen receptors (CARs) can re-direct T cells to target abnormal cells but their activity is limited by a profound defect in antigen sensitivity, the source of which remains unclear. Here we show that, while CARs have a >100-fold lower antigen sensitivity compared to the T cell receptor (TCR) when antigen is presented on antigen-presenting-cells (APCs), they have nearly identical sensitivity when antigen is presented as purified protein on artificial surfaces. We next measured the impact of engaging accessory receptors (CD2, LFA-1, CD28, CD27, 4-1BB) on antigen sensitivity by adding their purified ligands. Unexpectedly, we found that engaging CD2 or LFA-1 improved TCR antigen sensitivity by 125 and 22-fold, respectively, but only improved CAR sensitivity by <5-fold. This differential effect of CD2 and LFA-1 engagement on TCR versus CAR sensitivity was confirmed using APCs. We found that sensitivity to antigen can be partially restored by fusing the CAR variable domains to the TCR CD3*ε* subunit (also known as a TRuC), and fully restored by exchanging the CAR variable domains with the TCR*αβ* variable domains (also known as STAR or HIT). Importantly, these improvements in TRuC and STAR/HIT sensitivity can be predicted by their enhanced ability to exploit CD2 and LFA-1. These findings demonstrate that the CAR sensitivity defect is a result of their inefficient exploitation of accessory receptors, and suggest approaches to increase sensitivity.

## Introduction

Adoptive cell transfer (ACT) of genetically engineered T cells expressing Chimeric Antigen Receptors (CARs) is a clinically approved cancer therapy for haematological malignancies (1, 2). CARs are synthetic receptors that are typically generated by the fusion of an antibody-derived, antigen-binding single-chain variable fragment (scFv) with intracellular signalling motifs from the cytoplasmic tails of the T cell receptor (TCR) complex. Although administration of CAR-T cells targeting the surface antigens CD19, CD20, and B cell maturation antigen (BCMA or CD269) on malignant B cells results in an excellent initial response, patients often relapse when malignant cells emerge with reduced levels of target antigens (3–8). One likely explanation for this escape is that CARs require 100 to 1000-fold higher antigen densities to induce T cell activation compared to the native TCR (9–11). The mechanism underlying this profound defect in antigen sensitivity, which is seen with both proximal (10, 11) and distal readouts of T cell activation (9), remains unclear.

One approach to improving CAR function has focused on varying the stalk/hinge region and/or the cytoplasmic signalling domains. There are several commonly used hinges, including from CD8a, CD28, and IgG1. Most CARs use the cytoplasmic domain of the TCR *ζ*-chain for signalling, either alone (1^st^ generation) or in combination with the CD28 or 4-1BB cytoplasmic signalling domains (2^nd^ generation) (12–15). A study comparing the ability of several of these CARs to kill target cells with very low antigen densities found that the CARs that performed best had the CD28 hinge and the signalling domain from *ζ*-chain, either alone or in combination with the CD28 domain (16). Other studies have replaced the TCR *ζ*-chain with the cytoplasmic chain of the CD3*ε* subunit of the TCR/CD3 complex (11, 17, 18).

A second approach to improving CAR function has focused on exploiting all the signalling domains present in the TCR/CD3 complex. For example, eTruC receptors fuse the scFv directly to the extracellular domain of CD3*ε* (19) whereas STARs (also called HIT receptors) replace the variable domains of the TCR with the scFv variable domains (20, 21). Using a xenograft carcinoma model with EGFR as the target antigen a STAR outperformed an eTruC, and both outperformed CARs (20). The precise mechanisms underlying the these performance differences are unclear.

The TCR is known to have remarkable antigen sensitivity; it is able to recognise even a single peptide major-histocompatibility-complex (pMHC) on cells (22). Diverse mechanisms have been shown to contribute to this sensitivity (23). These include having multiple immunoreceptor tyrosine-based activation motifs (ITAMs) (24, 25), using the TCR co-receptors CD4 or CD8 (26, 27), and exploiting TCR accessory receptors such as LFA-1 (28) and CD2 (29). Despite the known importance of accessory receptors in enhancing TCR antigen sensitivity, their contribution to CAR antigen sensitivity has not been measured. Interestingly, CD2 has been shown to affect T cell activation by 1^st^ generation CARs but its impact on CAR antigen sensitivity is presented unknown (30).

Here, we take advantage of a shared pMHC antigen ligand to directly compared the antigen sensitivity of CARs with the native TCR. We show that, while CARs exhibit a >100-fold lower antigen sensitivity than TCRs to antigen presented on cells, they exhibit nearly identical sensitivities to antigen in the absence of accessory receptor ligands. We then demonstrate that engagement of accessory receptors only modestly increases the sensitivity of CARs to antigen, despite dramatically enhancing the sensitivity of the TCR. Finally, we show that TruCs and STARs/HITs have greater antigen sensitivity than CARs, and that this correlates with their ability to exploit CD2 to enhance this sensitivity. Our work helps explain the profound defect in CAR sensitivity and suggests ways to improve it for therapeutic purposes.

## Results

### Standard CAR designs exhibit reduced sensitivity compared to the TCR when antigen is presented on APCs but not when presented in isolation

To compare the antigen sensitivity of TCRs and CARs, we utilised the C9V variant (9V) of the NY-ESO-1_157–165_ cancer testis peptide antigen expressed on HLA-A*02:01, because it is recognised by both the 1G4 TCR (31, 32) and the D52N scFv (33) (Fig. 1A). While D52N binds to pMHC in a similar orientation to the 1G4 TCR (34), it binds 9V pMHC with a higher affinity (33). We produced five CAR designs by fusing the D52N scFv to either the CD28, CD8a, or IgG1 hinge coupled to either the TCR *ζ*-chain alone (1^st^ generation) or in combination with the CD28 signalling chain (2^nd^ generation) (Fig. S1).

**Figure 1:**
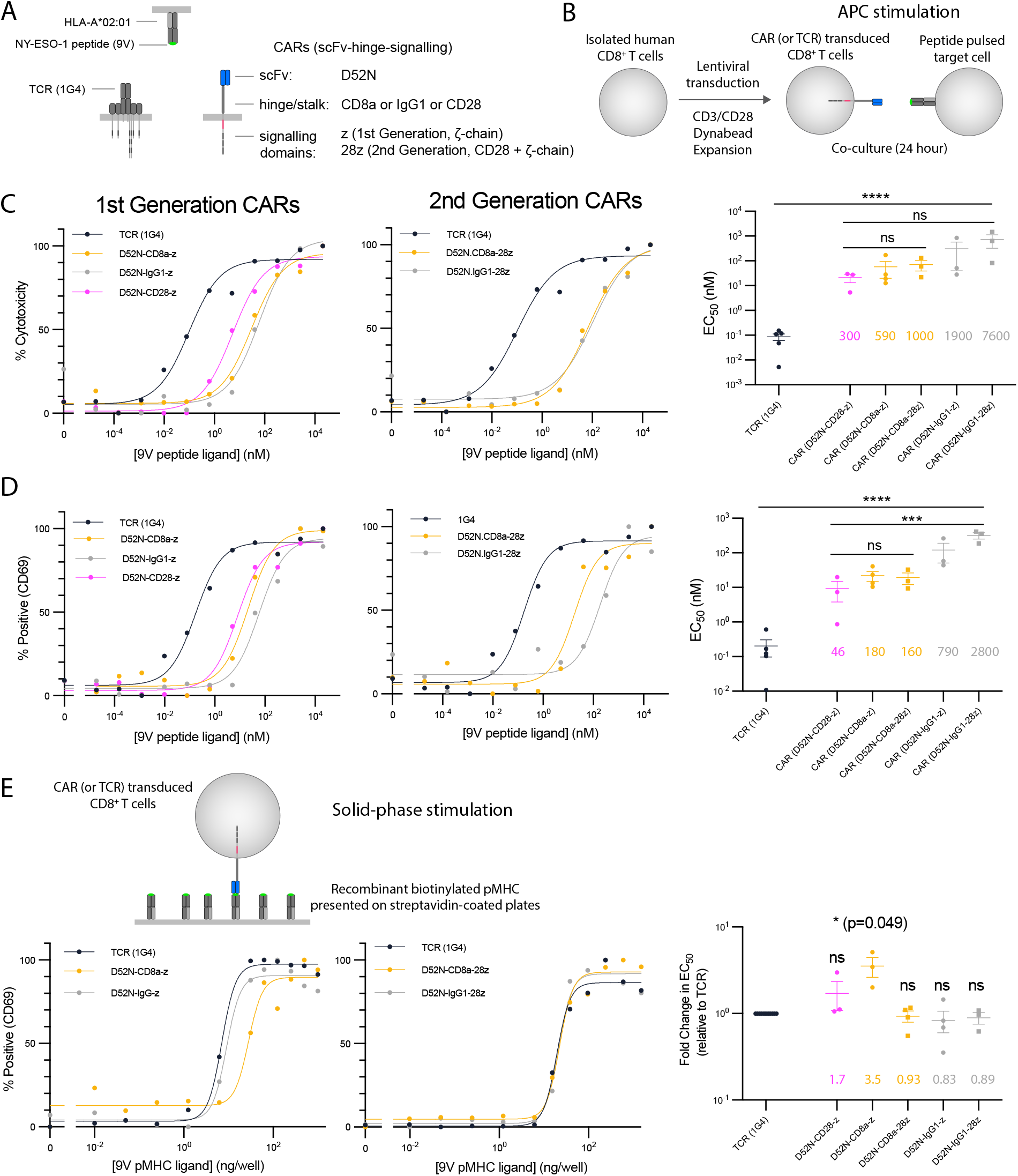
CARs show reduced sensitivity compared to the TCR when antigen is presented on APCs but not when presented as purified protein. **(A)** Schematic of antigen receptors. The 1G4 TCR and the D52N scFv both recognise the 9V NY-ESO-1 peptide antigen presented on HLA-A*02:01. CARs using the CD8a hinge contain the CD8a transmembrane domain whereas CARs using the IgG1 or CD28 hinges contain the CD28 transmembrane domain. **(B)** Schematic of APC stimulation system. **(C-D)** Representative dose-response showing **(C)** cytotoxicity by LDH release and **(D)** surface expression of CD69 for the TCR and the indicated CARs along with EC_50_ values from at least 3 independent experiments determined by fitting a Hill function to each dose-response curve. **(E)** Representative dose-response when purified biotinylated 9V pMHC ligand is presented on streptavidin-coated plates (left two plots) and EC_50_ values from at least 3 independent experiments (right). The EC_50_ values are compared using (C,D) one-way ANOVA or (E) one-sample t-test for a hypothetical mean of 1.0 on log-transformed values. Abbreviations: * = p-value≤0.05, ** = p-value≤0.01, *** = p-value≤0.001 **** = p-value≤0.0001.

Using a standard protocol similar to those employed in ACT (35), we transduced primary human CD8^+^ T cells with each antigen receptor and expanded them *in vitro* before co-culturing them with the HLA-A*02:01+ T2 target cell line pulsed with different concentration of antigen (Fig. 1B). We found that T cells expressing the 1G4 TCR were able to kill target cells (as measured by LDH release) at 300 to 7600-fold lower concentration of peptide antigen compared to CARs (Fig. 1C). We observed similar results when measuring the upregulation of the CD69 activation marker, albeit with lower 46 to 2800-fold changes (Fig. 1D). The large antigen sensitivity differences between the TCR and CARs could not readily be explained by receptor surface expression because the CARs were expressed at the same (or higher) levels than the TCR, as measured by pMHC tetramer binding (Fig. S2). This >100-fold higher sensitivity of the TCR is consistent with two previous reports (9, 10) that utilised different hinges and different signalling chains (2nd generation CARs with 4-1BB coupled to the *ζ*-chain). Our finding that a CAR with the CD28 hinge had the highest antigen sensitivity is also consistent with a previous report (16). Taken together, these results validate our antigen receptor system and suggest that reduced antigen sensitivity is a general feature of CARs.

We next compared antigen sensitivity of TCR and CARs when presented with plate immunobilized pMHC (Fig. 1E). This reductionist system allows precise control of TCR and accessory receptor ligands (32, 36–39). In striking contrast to the >100-fold difference in sensitivity when antigen was presented on cells, the TCR and CARs displayed similar antigen sensitivities when recognising purified antigen, with the largest difference being 3.5-fold (Fig. 1E).

### Ligands to the adhesion receptors CD2 and LFA-1 increase the antigen sensitivity difference between the TCR and CARs

Our finding that the ≳100-fold higher sensitivity of TCR compared to CARs is eliminated in a reductionist system provided an opportunity to explore the underlying mechanism. A key difference between cells and our reduced system is the presence of accessory receptor/ligand interactions involving T cell accessory receptors CD2, LFA-1, CD28, CD27, and 4-1BB (Fig. 2A). To investigate whether engagement of these receptors can account for the sensitivity differences, we tested their ability to increase antigen sensitivity by including, alongside pMHC, purified forms of their ligands at a concentration of (250 ng/well) previously shown to enhance T cell responses (32, 38, 40).

**Figure 2:**
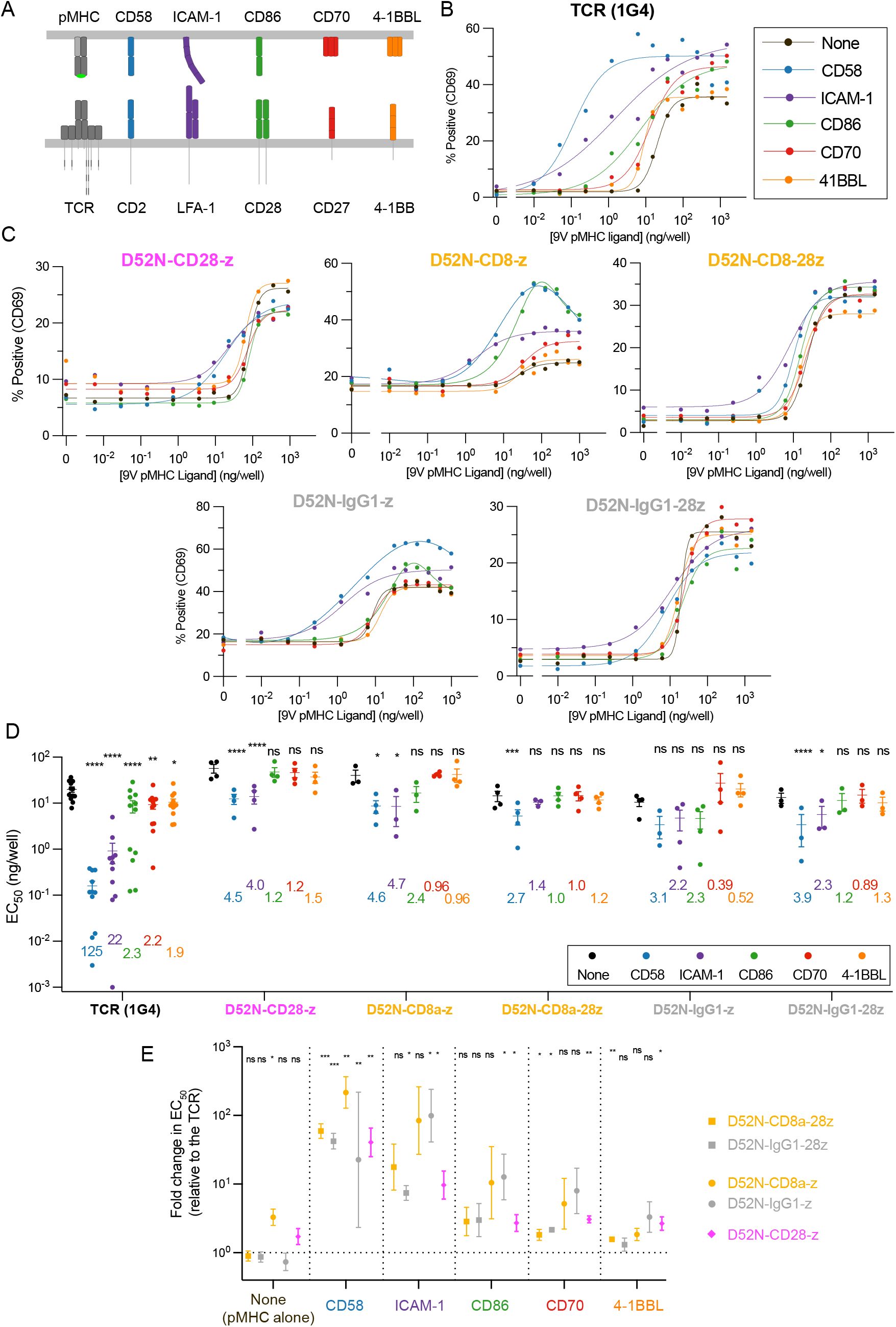
Systematic engagement of accessory receptors identifies that CARs are inefficient at exploiting the adhesion receptors CD2 and LFA-1 relative to the TCR. **(A)** Schematic of accessory receptors and their ligands. **(B-C)** Representative dose-response curves showing T cell activation by upregulation of surface CD69 measured by flow cytometry after 24 hours using the solid-phase stimulation assay. T cells were presented with purified pMHC alone (‘None’) or with a fixed concentration of 250 ng/well of the indicated accessory receptor ligand (colours) for the (B) TCR and (C) the indicated CARs. **(D)** The EC_50_ values for the indicated antigen receptor and purified ligand condition were obtained by fitting a Hill function to each dose-response curve. Individual EC_50_ values for each antigen receptor are from an independent experiment (N≥3). The numbers indicate the fold-change in EC_50_ induced by the accessory receptor ligand relative to pMHC alone (‘None’) and statistical significance is determined by a paired t-test on log-transformed data. **(E)** The data in (D) is presented in a different format showing the fold-change in EC_50_ between the TCR and the indicated CAR for pMHC alone or the indicated accessory receptor ligand. The fold-change is compared using a one-sample t-test to a hypothetical value of 0 on log-transformed data. Abbreviations: * =p-value≤0.05, ** = p-value≤0.01, *** = p-value≤0.001, **** = p-value≤0.0001

While ligands for CD2 (CD58), LFA-1 (ICAM-1), CD28 (CD86), CD27 (CD70), and 4-1BB (4-1BBL) all enhanced TCR antigen sensitivity, only CD58 and ICAM-1 increased CAR sensitivity (Fig. 2B-D). CD58 and ICAM-1 produced the largest increases in TCR antigen sensitivity (125- and 22-fold, respectively), while CD86, CD70, and 4-1BBL produced much smaller increases (Fig. 2D). Strikingly, CARs were much less efficient at exploiting these ligands than the TCR, with only CD58 and ICAM-1 increasing sensitivity, and only by 1.4 to 4.7 fold (Fig. 2D). When performing independent experiments (Fig. 2D, individual EC_50_ values), we isolated and transduced T cells from each donor with the TCR and one or more CARs. By always including the TCR, we coud express the antigen sensitivity of all CARs relative to the TCR (Fig. 2E). This confirmed that the TCR and CARs were similarly sensitive when stimulated with only pMHC, while the TCR was more sensitive when ligands to accessory receptors were present, with the largest differences observed when including ligands to CD2 or LFA-1.

To confirm these findings with another readout of T cell activation we measured production of the inflammatory cytokine IFN*γ*. As observed when using CD69 upregulation as a readout, accessory receptor ligands increased TCR sensitivity much more than CAR sensitivity, with CD2 ligands producing the biggest increases (Fig. S3).

It has previously been reported that tonic signalling by CARs can lead to T cell dysfunction/exhaustion by various mechanisms, including altering the expression of surface receptors (41–43), raising the possibility that tonic CAR signalling abrogates antigen sensitivity. To investigate this we first measure expression levels of accessory receptors (Fig. S4A) and exhaustion markers LAG-3, PD-1 and TIM-3 (Fig. S4B). These were indistinguishable between TCR and CAR-transduced T cells, except for a <2-fold increase in TIM-3. Next, we showed that transduction of a CAR did not affect the sensitivity of an orthogonal TCR recognising a viral peptide, with or without the CD2 ligand (Fig. S4C). This ruled out tonic signalling as an explanation for the defect on CAR antigen sensitivity.

Supra-physiological affinities can impair TCR signalling and reduce antigen sensitivity (37, 44, 45) and lowering the affinity of CARs has been shown to improve their *in vivo* activity (46). It follows that the higher affinity of the D52N scFv than the 1G4 TCR for the 9V (~50-fold higher at 37°C, Fig. S5A) could account for the defect in CAR sensitivity. To investigate this we identified a lower-affinity pMHC that bound the D52N scFv with the same affinity that the 1G4 TCR binds the 9V pMHC (Fig. S5A; 4A pMHC). When using these matched affinity pMHC antigens the difference in antigen sensitivity between the TCR- and CAR-transduced T cells was increased rather than decreased (Fig. S5B-E), demonstrating that the higher affinity of the CAR for antigen cannot account for its lower sensitivity.

The CD8 co-receptor binds pMHC, raising the possibility that it contributes to the difference in TCR and CAR sensitivity for pMHC antigen. To investigate this, we repeated the solid-phase stimulation assay using a pMHC variant with point mutations that abolish CD8 binding (47) (Fig. S6). Eliminating CD8 binding had no impact on the CAR sensitivity to pMHC and only a modest impact on antigen sensitivity to 9V pMHC. Interestingly, eliminating CD8 binding abolished TCR recognition of a very low-affinity 4A pMHC, consistent with previous work showing that CD8 has disproportionate impact on recognition of low-affinity antigens by TCR (48, 49). These findings show that CD8 binding does not account for the profound difference on CAR and TCR sensitivity.

In summary, the antigen sensitivities of the TCR and CARs are similar when presented with purified antigen in isolation, and antigen sensitivity of the TCR is enhanced far more than CARs by when including ligands for accessory receptors, especially CD2. This difference in TCR and CAR antigen sensitivity is not a result of differences in tonic-signalling, affinity for antigen, or the contribution of the CD8 co-receptor.

### Abrogating the CD2 and LFA-1 interaction reduces the antigen sensitivity difference between the TCR and CARs

Our results using an artificial system indicate that the antigen sensitivities of the TCR and CARs were similar when recognising purified antigen in isolation but exhibited large differences with the addition of purified ligands to CD2 or LFA-1 (Fig. 1–2). To investigate the role of these accessory adhesion receptor interactions in target cell recognition, we utilised the HLA-A*02:01 + U87 glioblastoma cell line, which expresses CD58 and ICAM-1 (Fig. S7A-B). We compared the TCR to the 1st generation CD8a hinge CAR (D52N-CD8a-z) because this CAR displayed the largest increase in antigen sensitivity when adding purified CD58 and ICAM-1 (Fig. 2D). We used blocking antibodies (Fig. 3A-D) or CRISPR (Fig. S7A-B, Fig. 3E-H) to abrogate CD58 and/or ICAM-1 engagement, and quantitated the effect on T cell sensitivity to pMHC antigen by measuring CD69 and 4-1BB expression. There was a profound ~ 100-fold difference in antigen sensitivity between TCR and CAR-transduced T cells, as shown above with T2 cell targets, which decreased to ~ 20-fold when abrogating both the CD2 and LFA-1 interaction (Fig. 3C-D,G-H, left panels). This decrease was mainly the result of a decrease in antigen sensitivity of the TCR (Fig. 3C-D,G-H, right panels). The fact that the antigen sensitivity of the TCR remained 20-fold higher than the CAR indicates that other mechanisms, including perhaps other ligand interactions, contribute to its higher sensitivity. In support of this, the U87 cell line expresses LFA-1 ligands other than ICAM-1 (Fig. S7C). In summary, our experiments with antigen presented on artificial surfaces or cells suggest that TCRs have higher antigen sensitivities than CARs because they exploit the accessory receptors such as CD2 and LFA-1 more efficiently.

**Figure 3:**
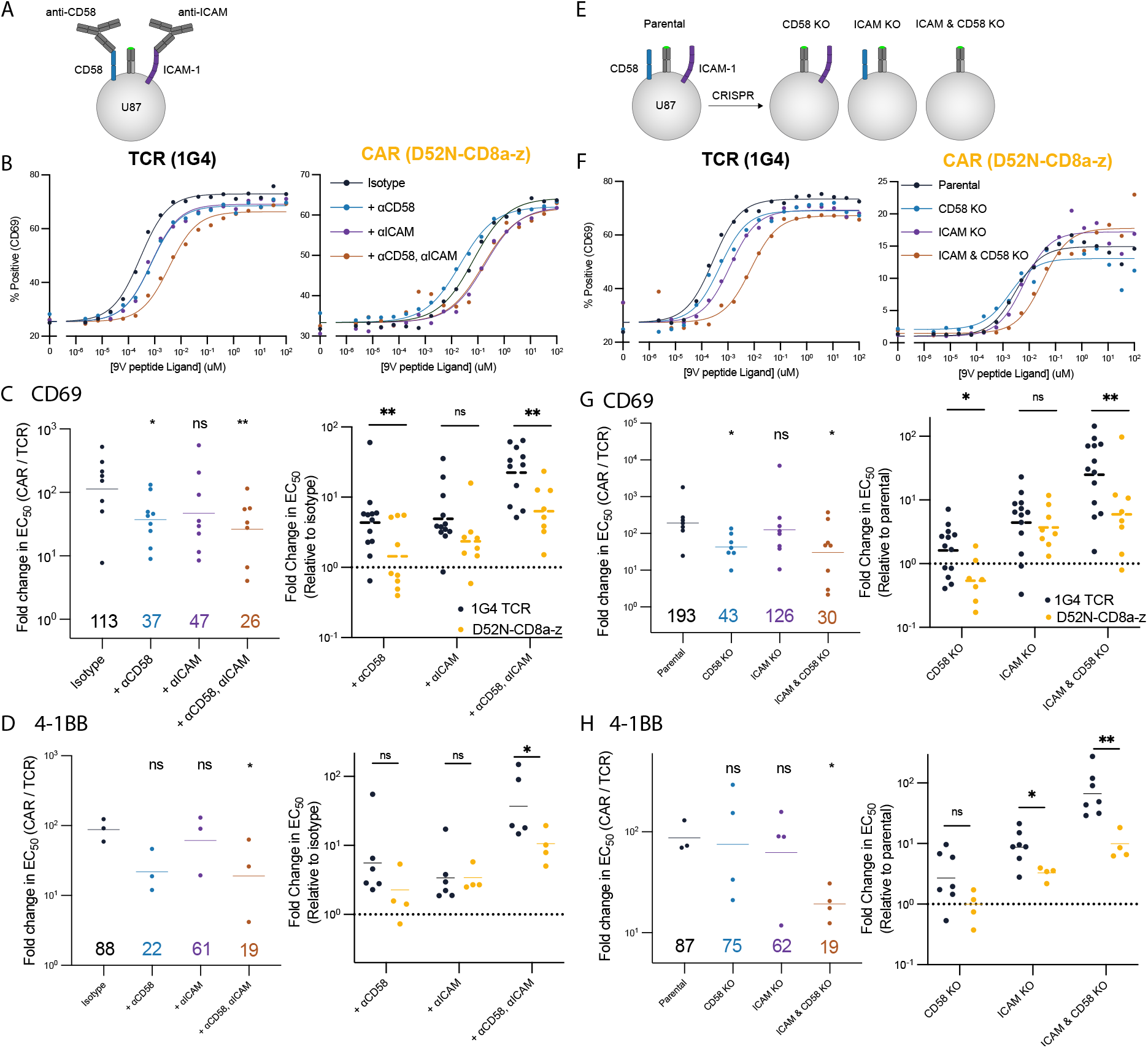
Abrogating the CD2 and LFA-1 adhesion interaction disproportionately impact the antigen sensitivity of the TCR compared to the CAR. **(A)** Schematic of CD58 and ICAM-1 blocking experiment on the HLA-A2+glioblastoma U87 target cell line. **(B)** Representative dose-response curves for the indicated blocking conditions for the TCR (left) and CAR (right). **(C-D)** Fold-change in EC_50_ between the CAR and TCR (left) or relative to the isotype (right) for (C) CD69 and (D) 4-1BB upregulation. **(E)** Schematic of CD58 and ICAM-1 knockout experiments. **(F)** Representative dose-response curves for the indicated target cell lines for the TCR (left) and CAR (right). **(G-H)** Fold-change in EC_50_ between the CAR and TCR (left) or relative to the isotype (right) for (G) CD69 and (H) 4-1BB. Individual EC_50_ values for CD69 or 4-1BB are determined by a fit to the dose-response curve from at least 3 independent experiments (each data point in C,D,G,H is from an independent experiment). The fold-change between the TCR and CAR is compared using a two-sample t-test to the isotype or parental line condition (left panel in C, D, G, H) or directly between the TCR and CAR (right panels in C, D, G, H) on log-transformed values. Abbreviations: * = p-value≤0.05, ** = p-value≤0.01, *** = p-value≤0.001, **** = p-value≤0.0001.

### STARs display TCR-like antigen sensitivity outperforming TRuCs and CARs by efficiently exploiting adhesion receptors

The ability of the TCR to exploit adhesion interactions has been shown to depend on both TCR signalling (50) and structural features of the TCR/pMHC interaction (23). The fact that conventional CARs lack signalling motifs present in the native TCR/CD3 complex has motivated the construction of new chimeric receptors. These include CARs containing cytoplasmic signalling chain of CD3*ε* (17, 18); TruCs, in which the scFv is fused to the extracellular domain of CD3*ε* and assembled into a complete TCR complex (19); and STARs, in which the TCR *α* and *β* chain variable domains are replaced with the antibody variable domains (20). These new chimeric receptors increasingly resemble the native TCR complex in terms of signalling components and structure (Fig. 4A).

**Figure 4:**
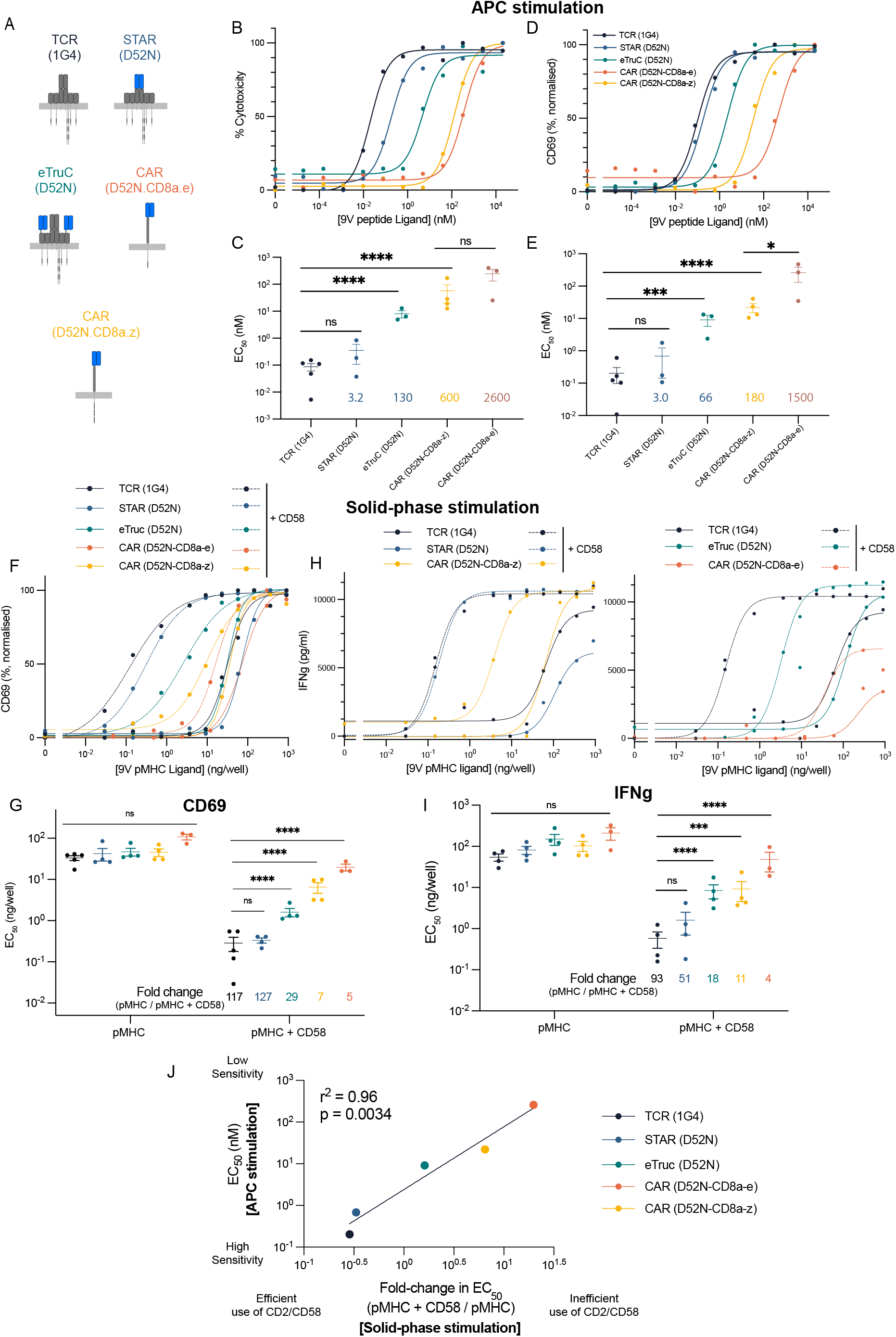
The ability of TCR-like chimeric antigen receptors to recapitulate the sensitivity of the TCR depends on the efficiency with which they are able to exploit the CD2 adhesion interaction. **(A)** Schematic of ‘TCR-like’ engineered antigen receptors. **(B-E)** T cells expressing the indicated antigen receptor were co-cultured with T2 target cells pulsed with different peptide antigen concentrations for 8 hours. Representative dose-response (top) and fitted EC_50_ values from at least 3 independent experiments (bottom) are shown for (B,C) cytotoxicity (measured by LDH release) and (D,E) CD69 upregulation. **(F-I)** T cells expressing the indicated antigen receptor were stimulated by a titration of purified pMHC alone (solid lines) or in combination with a fixed concentration of purified CD58 (dashed lines). Representative (F,H) dose-response curves and (G,I) fitted EC_50_ values from at least 3 independent experiments for (F,G) CD69 upregulation and (H,I) IFN*γ* production. **(J)** The averaged EC_50_ values for CD69 upregulation from the APC stimulation assay (from panel C) are plotted over the averaged fold-change in EC_50_ for CD69 induced by the addition of CD58 from the solid-phase stimulation assay (from panel G). The EC_50_ values are compared using a one-way ANOVA on log-transformed values (C,E,G,I). Abbreviations: * = p-value≤0.05, ** =p-value≤0.01, *** = p-value≤0.001, **** = p-value≤0.0001.

To directly compare the antigen sensitivities of these receptors using our system we generated versions containing the D52N variable domains (Fig. 4A). The CAR and eTruC incorporated the D52N scFv, which contains a linker between the variable domains, and bind its pMHC ligand (Fig. S5). The STAR incorporates the D52N variable domains into separate chains (Fig. S1). Because this lacks the linker present in the scFv we generated purified STAR and confirmed that it bound the pMHC, albeit with a 10-fold lower affinity than scFv (Fig. S8). When transduced into T cells the surface expression of these new chimeric receptors was indistinguishable from the 1G4 TCR (Fig. S2D, last three columns).

We next measured the sensitivity of these chimeric receptors to antigen presented on cells (APC stimulation) using target cell killing and CD69 upregulation as readouts (Fig. 4B-E). We found that the STAR performed identically to the TCR, while the eTruC was intermediate between them and the standard *ζ*-chain CAR. The *ε*-chain CAR was less sensitive than the *ζ*-chain CAR. To determine if adhesion interactions can account for these differences, we examined the impact of CD2 engagement on sensitivity to antigen presented on plates (solid-phase stimulation), using CD69 upregulation and cytokine production as read-outs (Fig. 4F-I). As before, we found nearly identical antigen sensitivities for all antigen receptors when presented with purified antigen alone. Addition of the CD2 ligand CD58 increased antigen sensitivity by different amounts, mirroring the antigen hierarchy observed with APC stimulation. Indeed, the efficiency with which an antigen receptor was able to exploit CD2 engagement directly predicted its antigen sensitivity measured using cells (Fig. 4J). We repeated the solid-phase stimulation assay using the LFA-1 ligand ICAM-1 and found a similar conclusion, albiet with lower fold-changes (Fig. SS9). Taken together, these results suggest that the antigen sensitivity of these TCR-like chimeric antigen receptors depends on their ability to exploit the CD2/CD58 and LFA-1/ICAM-1 adhesion interactions.

## Discussion

The ability of CAR T cells to control cancer cell mutants that express low antigen levels will depend on their sensitivity to antigen (1, 2, 6). We have shown here that several CAR formats, including 1^st^ and 2^nd^ generation CARs have a >100-fold lower sensitivity to antigen than the TCR. We further showed that this low sensitivity is the result of a failure of these CARs to efficiently exploit the adhesion receptors CD2 and, to a lesser extent, LFA-1. Finally, we show that this failure is reversed when chimeric receptors are redesigned to match more closely the native TCR structure.

Because it difficult to vary the CAR target antigen concentrations on cells, only a handful of studies have directly measured the antigen sensitivity of CARs. Consistent with our work, these studies reported a ~ 100-1000-fold defect in CAR antigen sensitivity compared to the TCR. Using a CAR containing the variable domains of a TCR, Harris et al (9) showed that both 1^st^ and 2^nd^ generation CARs exhibited a ~ 100-fold lower antigen sensitivity than the native TCR. Wang et al (51) found similar defects when using primary T cells and, consistent with our findings, observed only modest impacts of CD28 engagement on the antigen sensitivity of TCR and CARs (Fig. 2). Gudipati et al (10) report similar findings using antigens presented on planar bilayers, which contained ICAM-1, consistent with our result in our solid-phase stimulation system when ICAM-1 is included (Fig. 2E). When comparing the antigen sensitivities of different CARs, Majzner et al (16) found that CARs with the CD28 hinge produced the highest sensitivity. This was achieved with a 1st generation CAR and thus did not require signalling by CD28 or 4-1BB. We also found that the CD28 hinge CAR produced the highest antigen sensitivity among the CARs we tested (Fig. 1) and that CD28 or 4-1BB ligands produced only modest enhancements in antigen sensitivity (Fig. 2). Lastly, Salter et al (11) showed that incorporation of a proline rich region or the GRB2 SH2 domain into a 2nd generation CAR with 4-1BB signalling domain can enhance antigen sensitivity but these CARs continued to display lower antigen sensitivity than the 2nd generation CARs with CD28 signalling domains. We note that, while co-stimulation signals have a modest impact on antigen sensitivity, they are nevertheless critical for *in vivo* tumour control, presumably because they improving CAR-T cell persistence and increase cytokine production (52). Thus, our results are consistent with the previous studies, and extend them by identifying inefficient exploitation of adhesion receptors as a cause of the reduced antigen sensitivity of CARs.

While studies in mice suggested a modest role for CD2 (29, 53), it is clearly important in human T cells function (32, 54), including elimination of cancerous (55) and virus-infected (56) cells. Defects in CD58 (either loss of expression or mutations) have been reported in B cell and T cell lymphomas (57–59) and CD2 expression on tumour infiltrating T cells has been shown to correlate with their function in several cancers (60). Patients with B cell lymphomas with CD58 defects showed reduced progression-free survival when treated with axicabtagene ciloleucel CAR-T cell therapy (61). This implies that even the reduced ability of CARs to exploit CD2 can impact *in vivo* efficacy. Our finding that TruCs and STARs can more efficiently exploit CD2 to achieve higher antigen sensitivities is consistent with the finding that a STAR outperformed an eTruC, and that both outperformed CARs in an in vivo xenograft tumour model (20).

Although high antigen sensitivity is often beneficial, there are scenarios where low antigen sensitivity is desirable, such as when the target antigen is expressed at high levels on cancer cells and low levels on normal cells (62, 63). It has previously been shown that antigen sensitivity can be tuned by changing the affinity of the CAR (46, 63, 64) or by using transcriptional circuits (65). The results presented here show that antigen sensitivity can also be tuned by altering the CAR architecture. For example, the sensitivity hierarchy that we observe [STARS >TruCs >CAR (CD28 hinge) > CAR (CD8a hinge) > CAR (IgG1 hinge)] suggests that standard CARs may be preferred for targeting cancers that overexpress antigens also expressed on normal cells. In contrast, STARs may be preferred in cancers with low levels of target antigen or which commonly escape by reducing expression of the antigen. Importantly, STARs would remain susceptible to immune evasion by cancer cells losing expression of CD58 and/or ICAM-1. An advantage of tuning antigen sensitivity by changing the CAR architecture is that changes to the recognition domain are not required, reducing the risk of inadvertently altering its specificity.

The TCR, and indeed CARs, belong to a large and diverse group of surface receptors known as immunoreceptors or Non-catalytic Tyrosine-phosphorylated Receptors (NTRs) (66). The mechanism by which these receptors convert extracellular ligand binding into intracellular signaling, known as receptor triggering, remains debated. In the case of the TCR, allosteric conformational changes have been proposed as a triggering mechanism (67). While previous work has shown that grafting antibody variable domains to replace TCR variable domains produces a functional receptor (20, 68), it was unclear how this receptor compared to the native TCR. Our results here show that this chimeric receptor (STAR/HIT) is indistinguishable from the TCR in terms of antigen sensitivity. This observation is difficult to reconcile with allosteric models of TCR activation given the very limited conservation between antibody and TCR variable domains. Our results are, however, compatible with conformational changes induced by mechanical pulling forces.

In conclusion, we show that it is possible to engineer chimeric receptors with the same antigen sensitivities as the TCR, and that this requires that they efficiently exploit the adhesion receptors CD2 and LFA-1. This suggests a simple way to tune antigen sensitivity in order to optimise the functional effect of T cells. While our results suggest a strategy to reduce immune escape, it does not eliminate it because cancers can abolish expression of the target antigen. Other strategies, including targeting multiple antigens, may be necessary to further reduce escape (69). There is increasing interest in re-directing other immune cells, such as macrophages, using chimeric antigen receptors (70, 71). Since these cell do not usually express CD2, our work suggests that introducing CD2 or another adhesion receptor may be necessary to achieve the same remarkable antigen sensitivity as the TCR.

## Acknowledgements

We thank Marion H. Brown, Philipp Kruger, and John Nguyen for helpful discussion and reagents. We thank Linda Wooldridge and Christoph Renner for the DT227/8KA HLA-A2 and D52N CAR constructs, respectively.

## Materials & Methods

### Peptides

Peptides were synthesised at a purity of >95% (Peptide Protein Research, UK). 9V refers to a peptide derived from NY-ESO_157–165_ (`SLLMWITQV`), 4A is derived from the same sequence (`SLLAWITQV`), and SL9 refers to a peptide from HIV p17 GAG_77-85_ (`SLYNTVATL`).

### Protein production

HLA-A*02:01 heavy chain (UniProt residues 25–298) with a C-terminal BirA tag and β_2_-microglobulin were expressed as inclusion bodies in *E.coli*, refolded in *vitro* as described in (72) together with the relevant peptide variants, and purified using size-exclusion chromatography on a Superdex S75 column (GE Healthcare, USA) in HBS-EP buffer (10 mM M HEPES pH 7.4, 150 mM NaCl, 3 mM EDTA, 0.005% v/v Tween-20). Purified pMHC was biotinylated using the BirA enzyme (Avidity, USA).

His-tagged, soluble extracellular domain (ECD) of human CD58 was produced either in Freestyle 293F suspension cells (Thermo Fisher) or adherent HEK 293T cells. His-tagged, soluble versions of the ECD of human ICAM1, 41BBL, CD70 and CD86 were produced using adherent HEK 293T cells. Freestyle 293F suspension cells were transfected using Freestyle MAX reagent, as previously reported (32). Adherent HEK 293T cells were transfected using Roche X-tremeGENE HP transfection reagent following the manufacturer’s protocol. In both cases the resulting supernatant was filtered with a 0.45 μm filter and proteins were then purified using Ni-NTA agarose columns. Biotinylation was either performed *in vitro* after purification, or *in situ* by co-transfection (final proportion 10%) of a secreted BirA and adding 100 μM D-biotin to the growth media. Further purification and excess biotin removal was performed by size exclusion chromatography in HBS-EP.

D52N chains were produced as inclusion bodies in E. coli and refolded in vitro as described in (73), except that inclusion bodies were solubilised in 20 mM Tris-HCl (pH 8.0), 8 M urea, 2 mM DTT, refolding buffer contained 150 mM Tris-HCl (pH 8.0), 3 M urea, 200 mM Arg-HCl, 0.5 mM EDTA, 0.1 mM PMSF, and the refolding mixture was dialysed against 10 mM Tris-HCl (pH 8.5). The D52N dimer was purified on anion-exchange chromatography on a HiTrap Q column, followed by size-exclusion chromatography on a Superdex S200 column (both from GE Healthcare).

All purified proteins were aliquoted and stored at −80 °C until use.

### Lentiviral production

HEK 293T cells were seeded in DMEM supplemented with 10% FBS and 1% penicilin/streptomycin in 6-well plates to reach 60–80% confluency on the following day. Cells were transfected with 0.25 μg pRSV-Rev (Addgene, #12253), 0.53 μg pMDLg/pRRE (Addgene, #12251), 0.35 μg pMD2.G (Addgene, #12259), and 0.8 μg of transfer plasmid using 5.8 μl X-tremeGENE HP (Roche). Media was replaced after 16 hours and supernatant harvested after a further 24 hours by filtering through a 0.45 μm cellulose acetate filter. Supernatant from one well of a 6-well plate was used to transduce 1 million T cells.

### T cell production

Human CD8+ T cells were isolated from leukocyte cones purchased from the National Health Service’s (UK) Blood and Transplantation service. Isolation was performed using negative selection. Briefly, blood samples were incubated with Rosette-Sep Human CD8+ enrichment cocktail (Stemcell) at 150 μl/ml for 20 minutes. This was followed by a 3.1 fold dilution with PBS before layering on Ficoll Paque Plus (GE) at a 0.8:1.0 ficoll to sample ratio. Ficoll-Sample preparation was spun at 1200 x*g* for 20 minutes at room temperature. Buffy coats were collected, washed and isolated cells counted. Cells were resuspended in complete RMPI (RPMI supplemented with 10% v/v FBS, 100 Units/ml penicillin, 100 μg/ml streptomycin) with 50 U/ml of IL-2 (PeproTech) and CD3/CD28 Human T-activator Dynabeads (Thermo Fisher) at a 1:1 bead to cell ratio. At all times isolated human CD8+ T cells were cultured at 37 °C and 5% CO_2_.

1 million cells in 1 ml of media were subsequently transduced on the following day using lentivirus encoding for the various constructs (e.g., TCR), per the section on lentiviral transduction. On days 2 and 4 post-transduction, 1 ml of media was exchanged and IL-2 was added to a final concentration of 50 U/ml. Dynabeads were magnetically removed on day 5 post-transduction. T cells were further cultured at a density of 1 million cells/ml and supplemented with 50 U/ml IL-2 every other day. T cells were used between 10 and 16 days after transduction.

### APC stimulation (co-culture with T2 cells)

T2 cells were stained with 5 μM Tag-It Violet (BioLegend) following the manufacturer’s protocol and then 60000 cells were seeded in a volume of 100 μL per well in a V-bottom 96 well tissue culture plate. T2 cells were then incubated with 100 μL of peptide dilution prepared to the desired concentration in complete RPMI for 1 hour at 37 °C. T2 cells were then washed, resuspended in 100 μl of complete RPMI and transferred to a flat-bottom 96 well tissue culture plate.

Primary T cells were counted and re-suspended in fresh media such that there were 30000 receptor positive cells per 100 μl. This volume was then added to the T2 cells transferred previously.

As controls for the LDH assay additional wells were prepared in triplicate containing only 30000 T cells for each construct, or only 60000 T2 cells. Both with media to the same final volume as the co-cultured cells. Triplicate wells serving as volume correction and media controls were also prepared.

Plates were then spun at 50 xg for 2 minutes and incubated for 8 hours at 37 °C. After this period plates were spun again at 50 xg for 2 minutes and a fraction of supernatant was removed for assessing LDH release. LDH release was assessed using CyQUANT LDH Cytotoxicity Assay Kits (Thermo Fisher) following the manufacturers protocol. EDTA was added to the remaining supernatant (final concentration 2.5 μM) and cells were detached by pipetting.

Cells were stained for CD69 (Clone FN50, dilution 1:200) as well as with pMHC tetramers (dilution 1:500). Stained cells were either analysed immediately or fixed with 1% formaldehyde in PBS and analysed on the following day.

T cells were discriminated from T2 cells by the absence of Tag-It Violet stain. Single T cells were identified on the basis of size and subsequent analysis performed on this population.

### Solid-phase plate stimulation

Pierce Streptavidin Coated High Capacity 96 well plates (Thermo Fisher) were washed with PBS and dilutions of biotinylated pMHC in PBS were added to each well in a 50 μl volume and incubated for 90 minutes at room temperature. Subsequently, plates were washed again with PBS and biotinylated accessory molecules (CD58, ICAM-1, CD86, CD70, 41BBL) were added at a fixed dose of 250 ng/well in 50 μl. Plates were again incubated for 90 minutes and then washed with PBS.

T cells were counted, washed in media and 75000 cells in 200 μl were dispensed per well, Plates were spun for 2 minutes at 50 xg and then incubated for 24 hours at 37 °C. Following this incubation a portion of supernatant was removed and stored for performing ELISAs. EDTA was added to the remaining supernatant (final concentration 2.5 mM) and cells were detached by pipetting. Collected cells were stained for CD45 (Clone HI30, dilution 1:200), CD69 (Clone FN50, dilution 1:200), 4-1BB (Clone 4B4-1, dilution 1:200) and with tetrameric PE-conjugated pMHC. Cells were analysed either immediately or 1 day later, following fixation with 1% formaldehyde in PBS.

### Generating U87 knockout cell lines

U87 cells (a kind gift of Vincenzo Cerundolo) were used to generate genetic knockouts for CD58, ICAM1, or both using CRISPR Cas9 RNP transfection. To generate CD58 KO cells, 50,000 U87 cells were seeded in a 24-well plate and transfected the next day using Lipofectamine CRISPRMAX Cas9 Transfection agent (Thermo Fisher), annealed crRNA:tracrRNA (TrueGuide CRISPR758411_CR, GTCAATGCACAAGTTAGTGT, Thermo Fisher; A35506 for tracrRNA, Thermo Fisher), and TrueCut Cas9 Protein v2 (Thermo Fisher, A36496) according to manufacturer’s instructions. Cells were FAC sorted and this mixed population was used for all experiments. Sorted CD58 KO cells or WT U87 cells were used to generate CD58/ICAM1 double KO cells or ICAM1 KO cells, respectively using the same protocol as above. Specifically, cells were transfected with crRNA:tracrRNA (TrueGuide CRISPR845351_CR, GCTATTCAAACTGCCCTGAT, Thermo Fisher) and subsequently FAC sorted. Accutase (Biolegend 423201) was used to dissociate cells before screening or sorting with anti-CD58 (TS2/9, Invitrogen 12-0578-42) or anti-ICAM1 (HA58, Biolegend 353114) to prevent potential digestion of CD58 or ICAM1 by trypsin. All cell lines showed similar expression of HLA-A2 by flow cytometry (clone BB7.2, Biolegend 343306).

### APC stimulation (co-culture with U87 cells)

25000 U87 cells were seeded in a tissue culture treated flat-bottom 96 well plate and grown overnight. On the following day the media was removed from these cells and they were incubated with peptides prepared to the appropriate concentration in complete DMEM (DMEM supplemented with 10% v/v FBS, 100 Units/ml penicillin, 100 μg/ml streptomycin) for 1 hour at 37 °C.

If blocking antibodies were used then the appropriate amount of T cells were incubated for 30 minutes prior to addition to the U87 cells with either anti-IgG1 ϰ Isotype control (BioLegend, Clone MOPC-21), anti-CD58 (BioLegend, Clone TS2/9) or anti-ICAM1 (eBioscience, Clone HA58) at a concentration of 10 μg/ml. Alternatively, both anti-CD58 and anti-ICAM1 together at a concentration of 5 μg/ml each (total antibody concentration 10 μg/ml).

Peptide containing media was then removed and 50,000 T cells per well were added. The co-culture was then spun for 2 minutes at 50 x*g*, and incubated for 4 hours at 37 °C. After this period a fraction of supernatant was removed for cytokine ELISAs and stored at −20 °C. EDTA was added to the remaining supernatant (final concentration 2.5 μM) and cells were detached by pipetting.

Cells were stained in PBS 1% BSA for CD45 (Clone HI30, dilution 1:200), CD69 (Clone FN50, dilution 1:200) and 4-1BB (Clone 4B4-1, dilution 1:200) as well as with PE-conjugated tetrameric pMHC (dilution 1:500). Stained cells were either analysed immediately or fixed with 1% formaldehyde in PBS and analysed on the following day.

T cells were discriminated from U87 cells by CD45 staining and/or an assessment of size and complexity. Single T cells were identified on the basis of size and subsequent analysis performed on this population.

### Flow cytometry

Tetramers were produced using refolded monomeric biotinylated pMHC and streptavidin-PE (Biolegend) at a 1:4 molar ratio. Streptavidin-PE was added in 10 steps with a 10 minute incubation at room temperature between each addition. 0.05–0.1% sodium azide was added for preservation and tetramers were kept for up to 3 months at 4 °C.

Samples were analysed using a BD LSR Fortessa X-20 (BD Biosciences) or CytoFLEX LX (Beckman Coulter) flow cytometer and data analysis was performed using FlowJo v10 (BD Biosciences).

### Electroporation of 868 TCR

868 TCR alpha and beta chains were amplified using PCR, adding a T7 promoter at the 5′ end. The resulting PCR product was ‘cleaned up’ using a NucleoSpin Gel and PCR clean-up kit (Macherey-Nagel). Capped and Poly(A) tailed mRNA was produced from this PCR product using a mMESSAGE mMACHINE™ T7 ULTRA Transcription Kit (ThermoFisher). mRNA was collected by lithium chloride precipitation, quality checked by gel electrophoresis and stored in single use aliquots at −80 °C.

For electroporation, T cells are collected and washed 3x with Opti-MEM (Gibco) and resuspended at a concentration of 25 × 10^6^ cells/ml. 5 × 10^6^ cells with 2 μg per million cells of each of the RNA for the TCRα, β and ζ chains. Cells were then aliquoted in 200 μl into an electroporation cuvette (Cuvette Plus 2mm gap BTX). Electroporation is performed using an ECM 830 Square Wave electroporation system (BTX) at 300 V for 2 ms. Cells are then transferred to pre-warmed complete RPMI at a density of 1 × 10^6^ cells/ml. Electroporated cells are used in assays 24 hours later.

### Immobilisation Assay

Following a plate stimulation assay, after cells were collected, plates were washed 3 times with PBS 0.05% TWEEN 20 (‘PBST’) and then stained with anti-HLA-A,B,C (clone W6/32, dilution 1:1000) in PBS for 2 hours at room temperature. Plates were then washed 3x with PBST and stained with secondary goat antimouse IgG IRDye 800CW (LI-COR) in PBS for a further 2 hours. Finally plates were washed one more time with PBST and then imaged using a LICOR Odyssey Sa (LI-COR). Integrated intensity per well is reported.

### ELISAs

Invitrogen Uncoated ELISA kits for IFNγ (Thermo Fisher) were used following the manufacturer’s protocol. Supernatants were either used immediately for ELISAs post-harvesting or stored at −20 °C for up-to 2 weeks. Supernatants were diluted using an empirically determined ratio before use in an ELISA so that quantities of assessed cytokines fell within the linear range of the kits.

### Surface Plasmon Resonance

D52N–pMHC interactions were analysed on a Biacore T200 instrument (GE Healthcare Life Sciences) at 37°C and a flow rate of 30 μl/min. Running buffer was HBS-EP (10 mM HEPES pH 7.4, 150 mM NaCl, 3 mM EDTA, 0.005% v/v Tween-20). Streptavidin was coupled to CM5 sensor chips using an amino coupling kit (GE Healthcare Life Sciences) to near saturation, typically 10,000–12,000 response units (RU). Biotinylated pMHCs (47 kDa) were injected into the experimental flow cells (FCs) for different lengths of time to produce desired immobilisation levels (300–1000 RU). FC1 and FC3 were used as reference FCs for FC2 and FC4, respectively. Biotinylated ECD of CD58 (24 kDa + 25 kDa glycosylation) was immobilised in the reference FCs at levels matching those of pMHCs. Excess streptavidin was blocked with two 40 s (D52N STAR) or 60 s (D52N scFv) injections of 250 μM biotin (Avidity). Before injections of purified D52N, the chip surface was conditioned with eight injections of the running buffer. Dilution series of D52N were injected simultaneously in all FCs starting from the lowest concentration, which was injected again after the highest concentration to confirm stability of pMHC on the chip surface. The duration of injections (20 or 180 s) was the same for conditioning and D52N injections. After every 2 or 3 D52N injections, buffer was injected to generate data for double referencing. In addition to subtracting the signal from the reference FC (single referencing), all D52N binding data were double referenced versus the average of the closest buffer injections before and after D52N injection to correct for small differences in signal between flow cells. D52N binding versus D52N concentration was fitted with the following model: 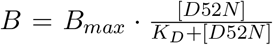, where *B* is the response (binding) and *B_max_* is the maximal binding.

### Sequences

D52N scFvs with the following sequence were produced by Absolute Antibody Ltd.

#### D52N scFv

~~~
EVQLLESGGGLVQPGGSLRLSCAASGFTFSTYQMSWVRQAPGKGLEW
VSGIVSSGGSTAYADSVKGRFTISRDNSKNTLYLQMNSLRAEDTAVY
YCAGELLPYYGMDVWGQGTTVTVSSAKTTPKLEEGEFSEARVQSELT
QPRSVSGSPGQSVTISCTGTERDVGGYNYVSWYQQHPGKAPKLIIHN
VIERSSGVPDRFSGSKSGNTASLTISGLQAEDEADYYCWSFAGGYYV
FGTGTDVTVLG
~~~

The D52N-IgG1 CARs contain a ‘HNG spacer sequence’ derived from the IgG1 hinge region, described in (74), and spliced with a spacer region from the CH2-CH3 regions of IgG1 as described in (75).

#### HNG Spacer

~~~
DPAEPKSPDKTHTCPPCP
~~~

The 1G4 TCR α and β chains are joined by a P2A linker peptide with an additional spacer and furin cleavage site, as described in (76). The sequence is given below.

#### Furin-P2A

~~~
GSRAKRSGSGATNFSLLKQAGDVEENPGP
~~~

### Independent experiments and data analysis

To produce independent measurements of EC_50_ (individual data points in figure panels) for a given antigen receptor, we produced a new batch of lentivirus which was used to transduce T cells isolated from a new leukocyte cone, which is provided by the National Health Service (NHS) in the UK and is obtained from human blood donors.

In each independent experiment, we included the TCR and one or more CARs to be tested and used pMHC antigen tetramers to evaluate the percent of T cells expressing each antigen receptor (Fig. S2C, transduction efficiency) and the surface level (gMFI of T cells expressing the antigen receptor) for each antigen receptor relative to the TCR (Fig. S2D). Although we observed variations in the transduction efficiency, the surface level of each antigen receptor was always at the same level or higher compared to the TCR.

As a result of differences in the transduction efficiency, we observed differences in the maximum number of T cells that could upregulate CD69 or 4-1BB across independent experiments for the same antigen receptor or across different antigen receptors (see y-axis in Fig. 2B,C for example). These differences reflect the percent of T cells that express the antigen receptor and can therefore respond to the presented antigen. Importantly, our study focused on measuring antigen sensitivity (EC_50_), which is defined as the concentration of antigen required to elicit half-maximal response. Therefore, variations in the maximum number of T cells able to respond are taken into account when measuring an EC_50_.

Statistical analysis was performed using Prism (GraphPad Software) or Excel (Microsoft). Curve fitting was performed using the robust nonlinear regression function in Prism or MATLAB (MathWorks) and the *EC*_50_ extracted from the fitted curves. Data was excluded from analysis if the computed fit was reported as ‘ambiguous’ in Prism, if the fit did not converge in 1000 iterations, or if the computed *EC*_50_ was outside of the tested ligand concentration.

## Supplementary Information

**Figure S1:**
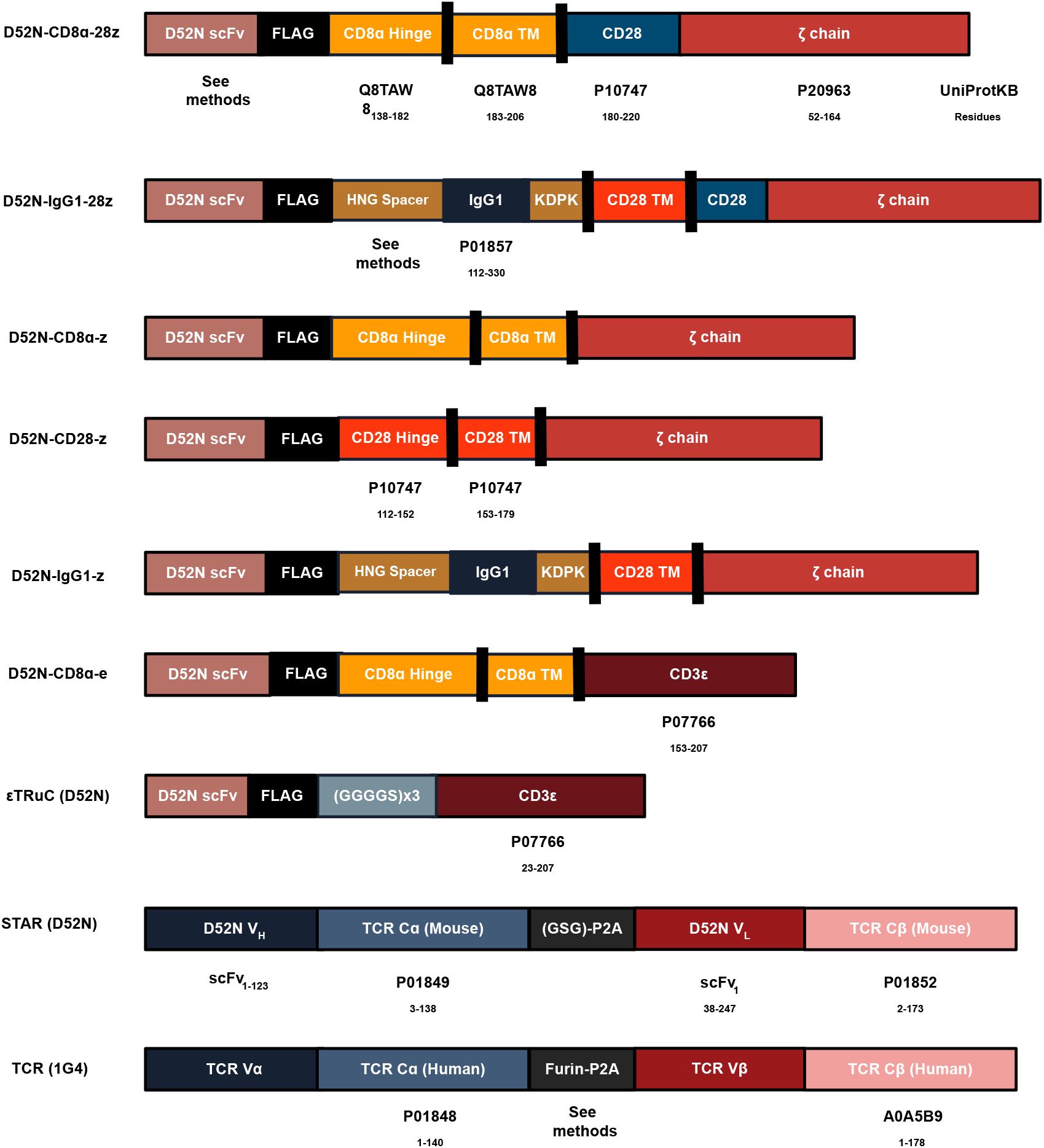
Schematic of antigen receptor architectures used in the study.

**Figure S2:**
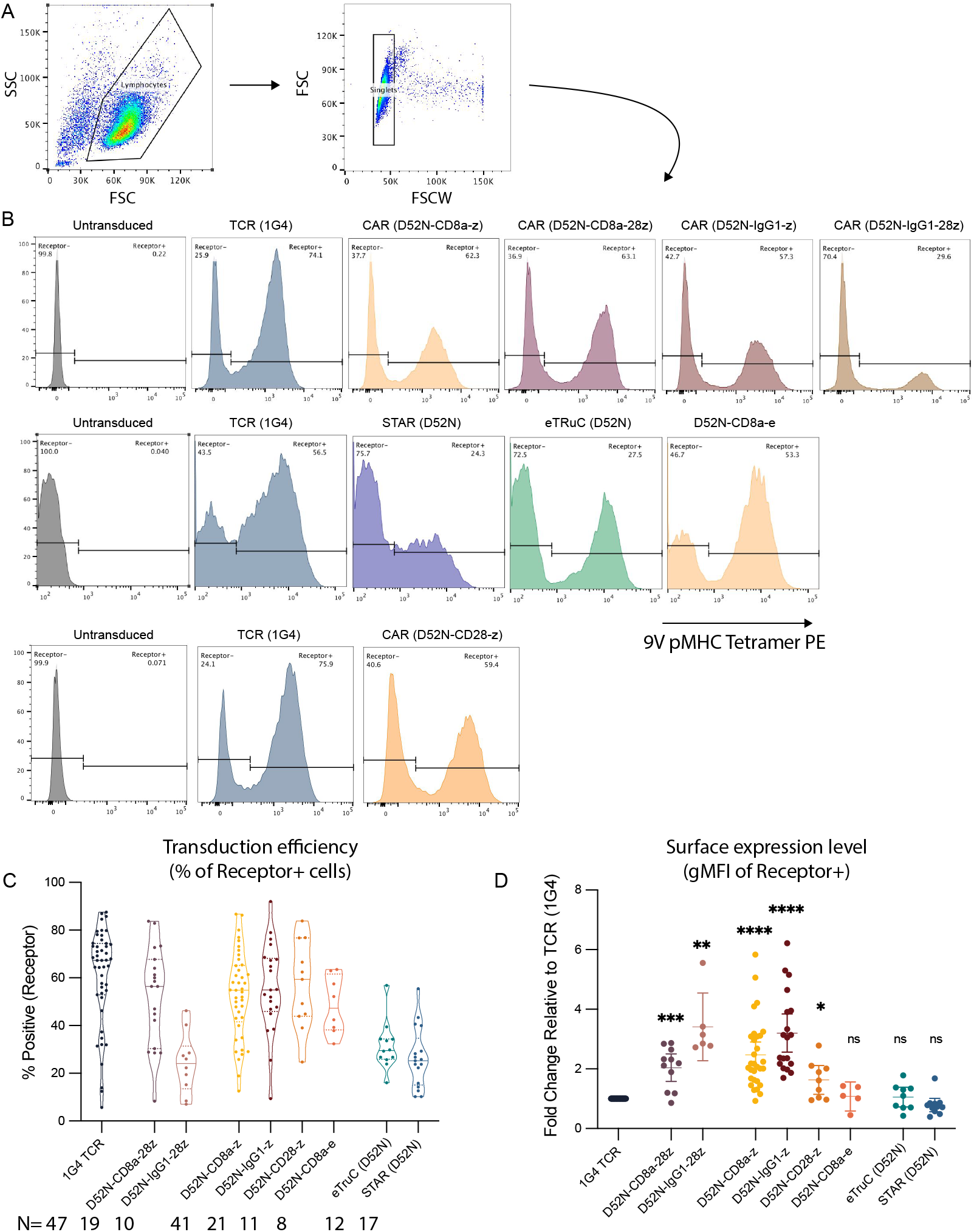
Surface expression of chimeric receptors was similar or higher compared to the TCR. **(A)** Gating strategy to identify single lymphocytes. **(B)** Representative flow cytometry histograms showing surface expression of the indicated surface receptor using fluorescent 9V pMHC tetramers. Untransduced T cells are used to determine the negative gate. Each row is an independent experiment and each column is the indicated antigen receptor with the TCR being included in all experiments performed in the study. **(C)** The percent of T cells expressing the indicated receptor (i.e. within receptor positive gate). **(D)** The fold-change in the surface expression of each chimeric antigen receptor relative to the TCR determined by the gMFI of T cells in the receptor positive gate. The surface expression of each antigen receptor was determined for every experiment carried out in the study and is shown in aggregate in panel C and D. Individual data points for each antigen receptors represents an independent experiment (N is shown below the labels in panel C), which is generated by producing lentivirus and transducing a new sample of primary human T cells (see Methods). The TCR contains the largest number of experiments because it was included in every experiment. A one-sample t-test is used to compare each chimeric receptor to the expression of the TCR (1.0) on log-transformed values. Abbreviations: * = p-value≤0.05, ** = p-value≤0.01, *** = p-value≤0.001, **** = p-value≤0.0001.

**Figure S3:**
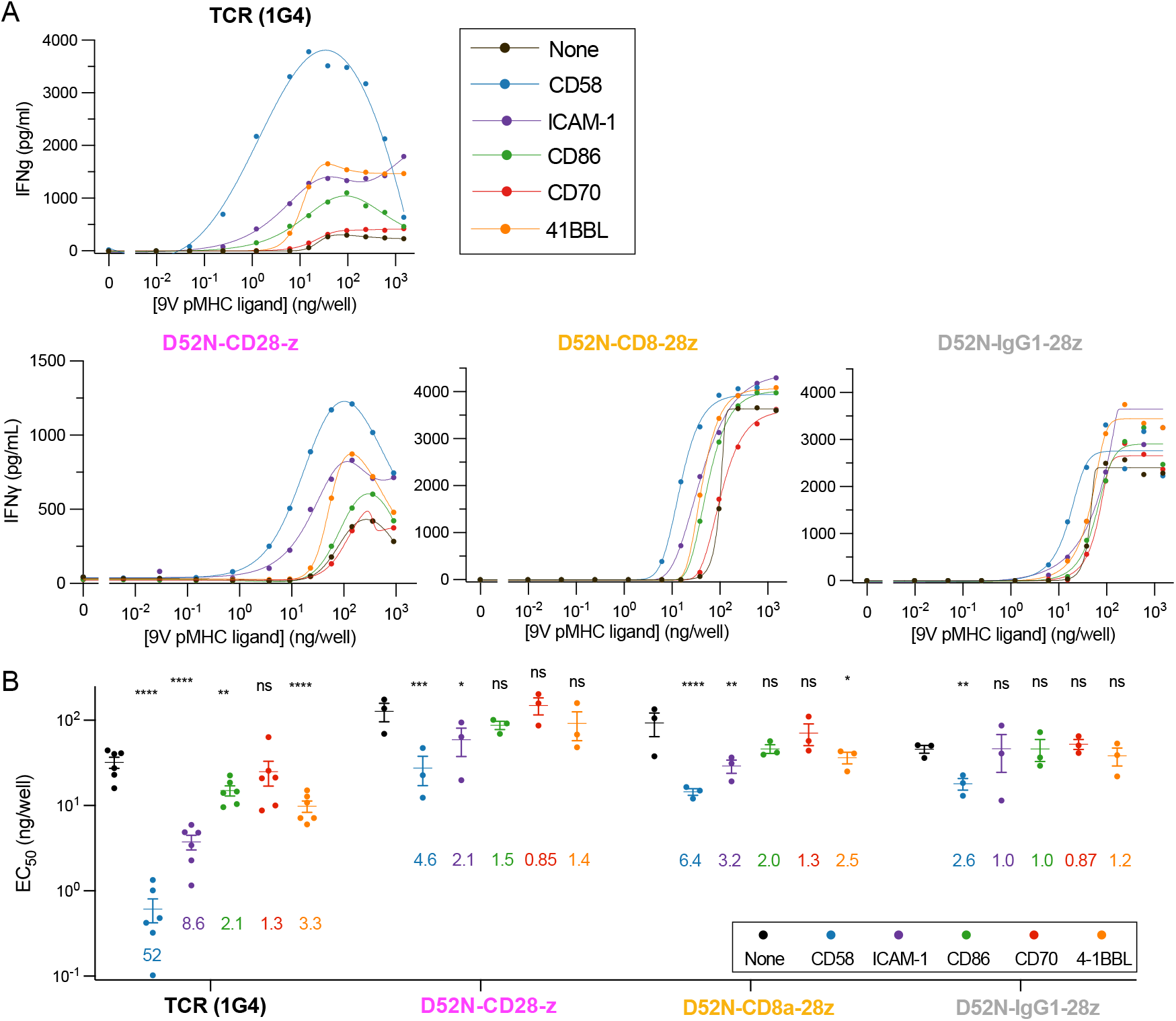
Extended data for Figure 2 confirming that CARs are inefficient at exploiting adhesion receptors when measuring IFN*γ* production. **(A)** Representative dose-responses using T cells expressing the indicated antigen receptor stimulated by a titration of purified pMHC alone (‘None’) or in combination with a fixed concentration of the indicated purified accessory receptor ligand (colours). The supernatant concentration of IFN*γ* was determined after 24 hours using ELISA. Each dose-response curve was fitted to obtain the EC_50_ and E_max_ values. **(B)** The EC_50_ values for the indicated antigen receptor and purified ligand. The coloured numbers indicate the fold-change in EC_50_ induced by the addition of the indicated accessory receptor ligand relative to pMHC alone (‘None’). The EC_50_ values for each ligand are compared to the ‘None’ condition using a paired t-test on log-transformed data. **(C)** The fold-change in E_max_ relative to pMHC alone (‘None’) for each antigen receptor and purified ligand. The fold-change is compared using a one-sample t-test to a hypothetical value of 0 on log-transformed data. Abbreviations: * = p-value≤0.05, ** = p-value≤0.01, *** = p-value≤0.001, **** = p-value≤0.0001.

**Figure S4:**
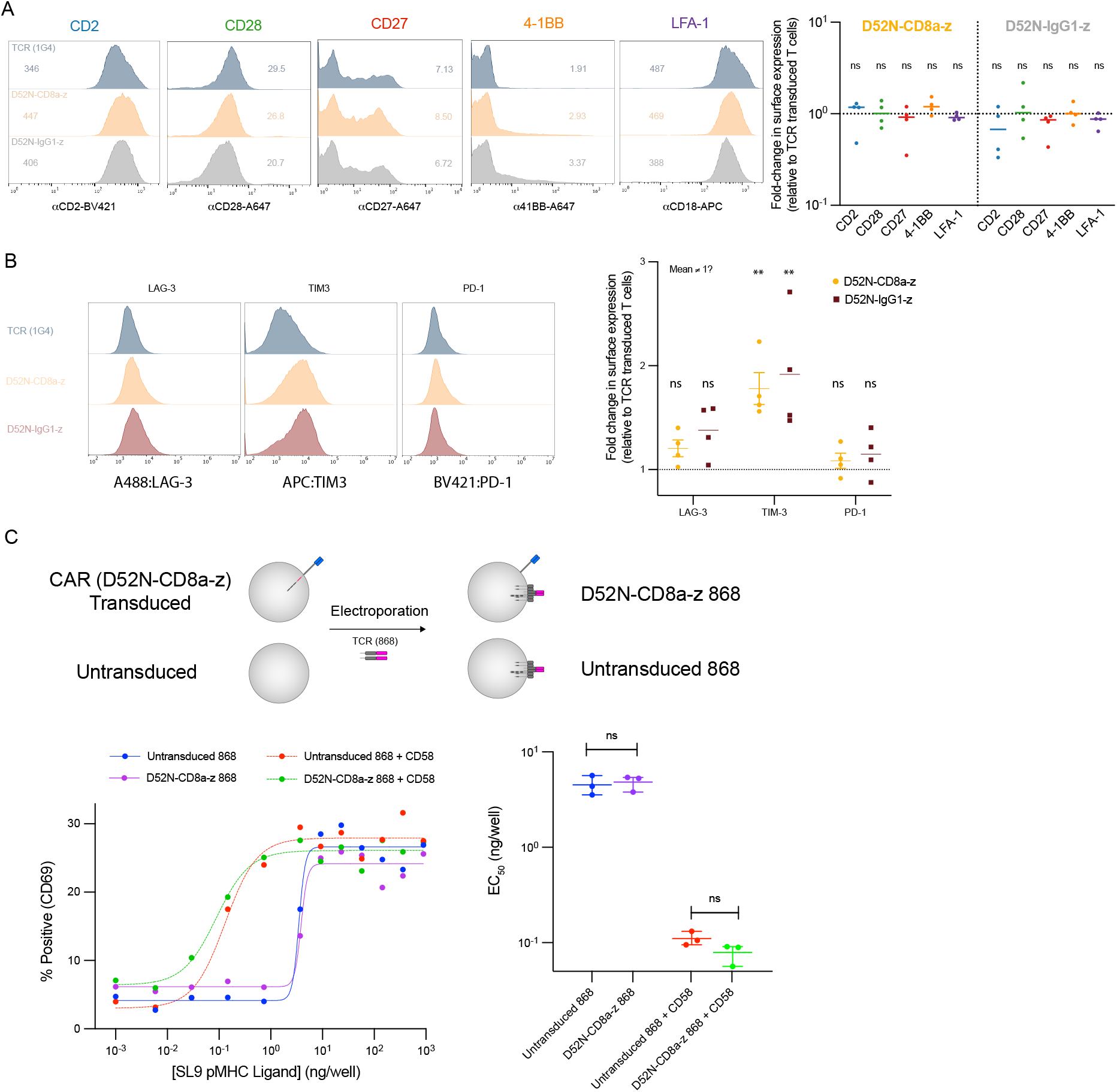
The CAR antigen sensitivity defect is not a result of exhaustion induced by tonic signalling. **(A)** Surface expression of the indicated co-stimulation receptors on T cells transduced with the TCR or the indicated CAR. Representative flow cytometry histograms (top) and fold-change across independent experiments (N=4, bottom). **(B)** Surface expression of the indicated co-inhibitory receptors on T cells transduced with the TCR or with the indicated CARs. Representative flow cytometry histograms (left) and fold-change across independent experiments (N=4, right). **(C)** CAR transduced or untransduced T cells are electroporated with the 868 TCR specific to the SL9 peptide antigen before being stimulated by purified SL9 pMHC with or without CD58. Representative dose-response (bottom left) and EC_50_ values across independent experiments (N=3, bottom right). A one-sample t-test is used to obtain a p-value for the null hypothesis that the indicated surface receptor expression differs from 1.0 on log-transformed values (panel A and panel B, right) and a two-sample t-test is used to compare log-transformed *EC*_50_ (panel C, bottom right). Abbreviations: * = p-value≤0.05, ** = p-value≤0.01, *** = p-value≤0.001, **** = p-value≤0.0001.

**Figure S5:**
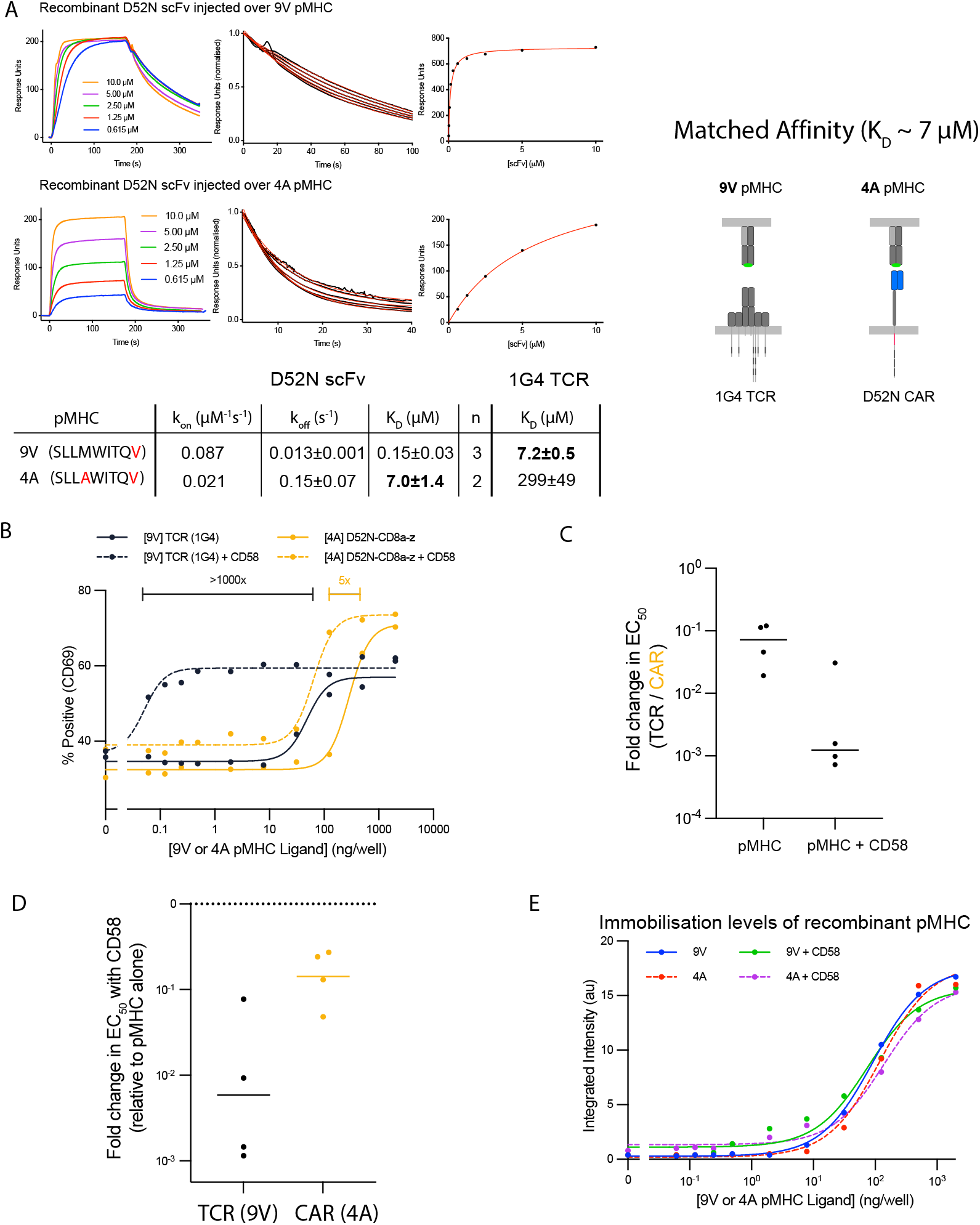
Matching the antigen affinity of the TCR and CAR increases the antigen sensitivity defect of CARs. **(A)** Binding between purified D52N scFv and the 9V (top) or 4A (bottom) pMHCs measured by surface plasmon resonance (SPR) showing the full sensogram (left), dissociation phase (middle) and steady state binding response (right). The kinetic *k*_off_ and equilibrium K_D_ are obtained by fitting the dissociation phase and steady-state binding response, respectively (red line is model fit). The kinetic *k*_on_ is derived from K_D_ and *k*_off_. The K_D_ values for the TCR are obtained from previous work (32). **(B)** Representative doseresponse for the TCR recognising the 9V pMHC and for the CAR recognising the 4A pMHC with (dashed line) or without (solid line) purified CD58 (250 ng/well). **C-D** Fold-change in EC_50_ between (C) the TCR recognising 9V and the CAR recognising 4A and (D) induced by the addition of CD58 for the indicated pMHC and antigen receptor across independent experiments (N=4). **(E)** Levels of presented pMHC for each condition as detected by the conformationally sensitive W6/32 antibody.

**Figure S6:**
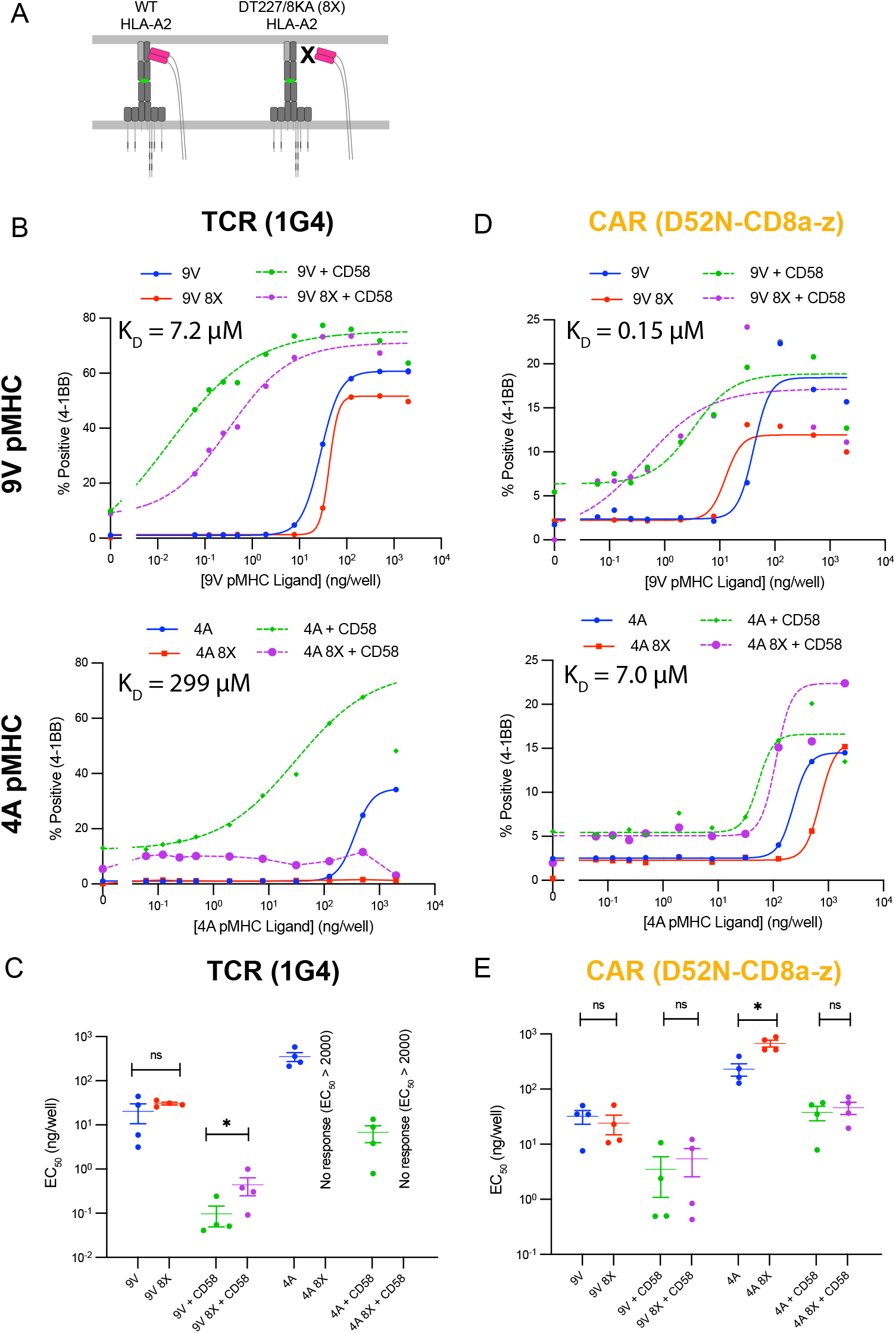
The CAR antigen sensitivity defect is independent of the CD8 co-receptor. **(A)** The DT227/8KA mutations in the HLA-A2 heavy-chain prevent binding by the CD8 co-receptor (referred to as 8X). **(B-E)** Representative dose-response curves (B,D) and summary measures across independent experiments (N=4) (C,E) for the 9V and 4A pMHC variants with or without CD58 for the TCR (B,C) and the CAR (D,E). A t-test is used to compare the EC_50_ values on log-transformed data. Abbreviations: * = p-value≤0.05, ** = p-value≤0.01, *** = p-value≤0.001, **** = p-value≤0.0001.

**Figure S7:**
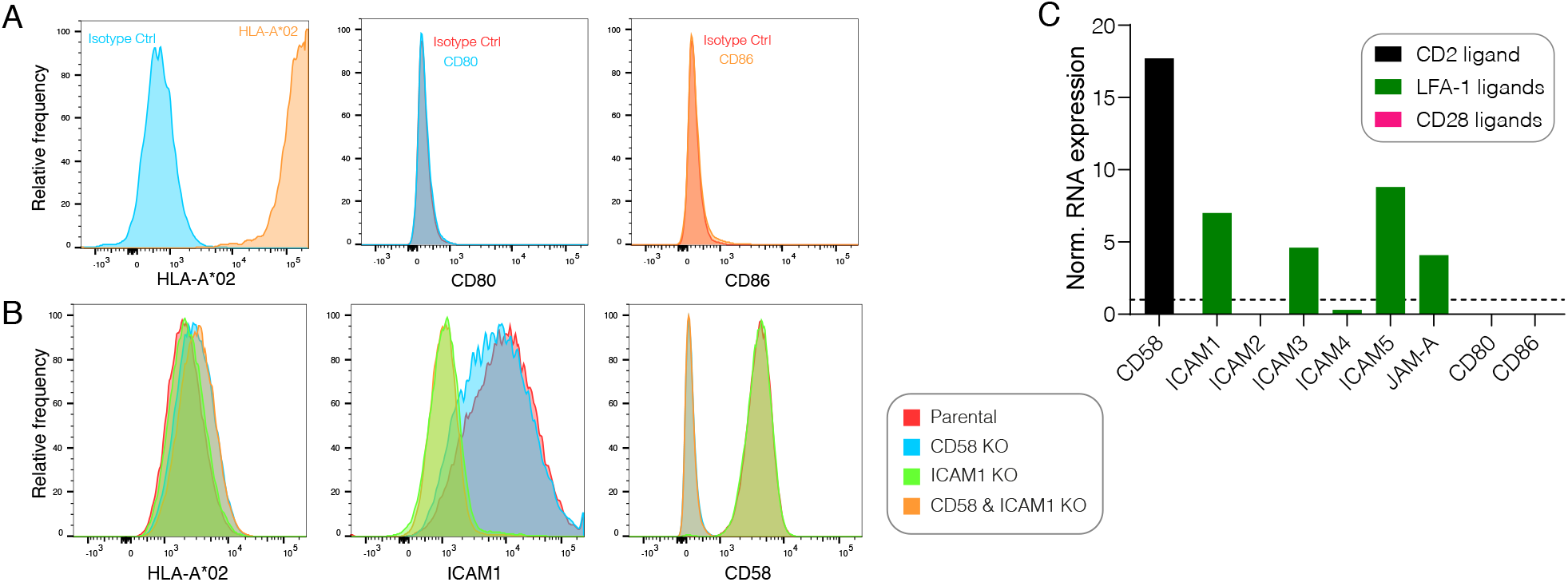
Expression of accessory receptor ligands on the U87 glioblastoma cell line. **(A)** Expression of HLA-A*02 (left), CD80 (middle), and CD86 (right) on parental U87 cells. **(B)** Expression of HLA-A*02 (left), ICAM-1 (middle), and CD58 (right) on the parental U87 cells or the indicated knockout cell lines. **(C)** Expression of the indicated molecule by RNA as reported in the Human Cell Atlas (77).

**Figure S8:**
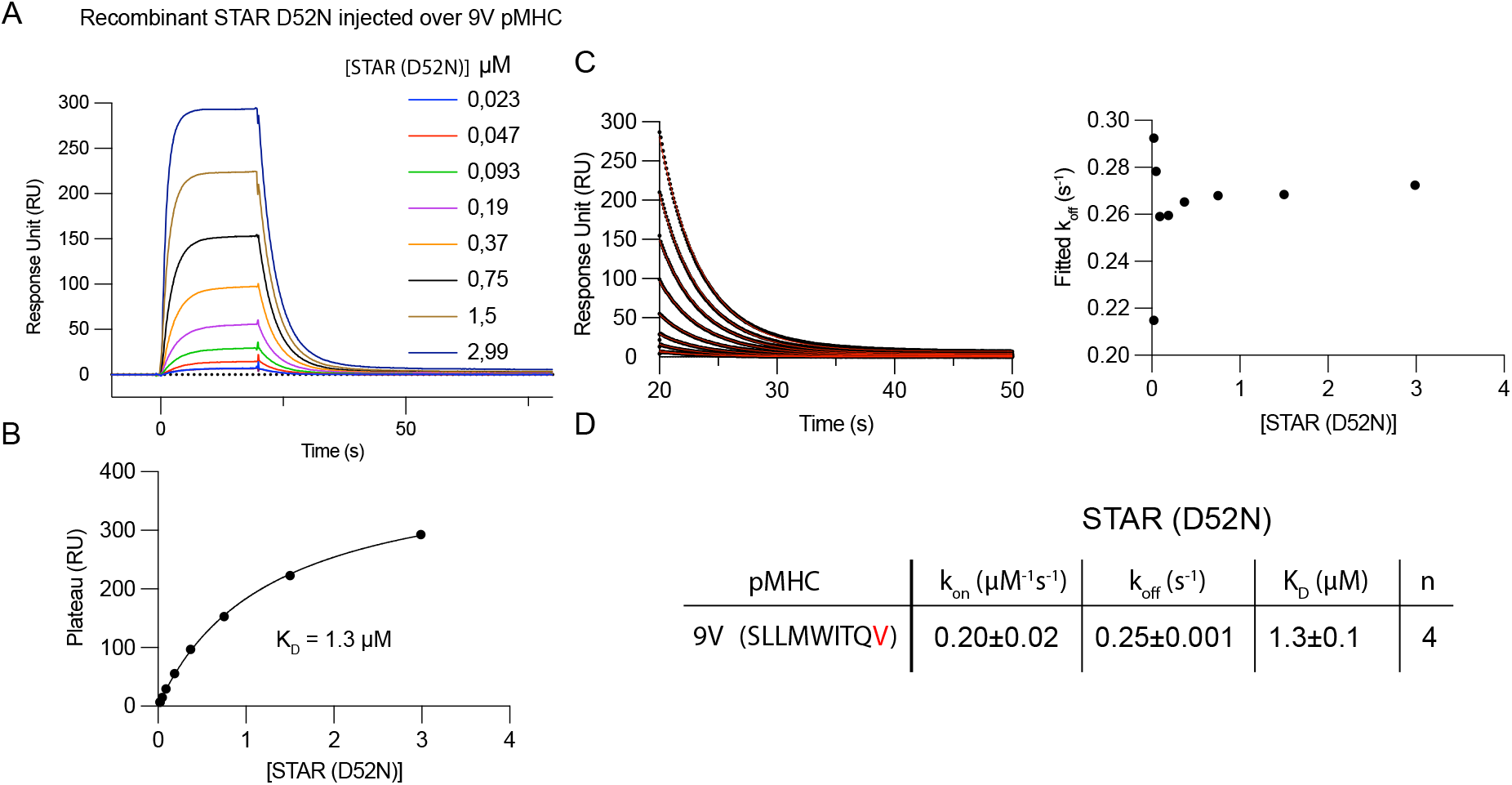
The binding affinity and kinetics between recombinant STAR (D52N) and 9V pMHC. **(A)** Representative binding between purified D52N STAR at the indicated concentration injected over a surface with immbolised 9V pMHC measured by surface plasmon resonance (SPR) showing the full sensogram. **(B)** The equilibrium K_D_ is obtained by fitting the steady-state response. **(C)** The kinetic *k*_off_ for each experiment is obtained by fitting the dissociation phase (left) and averaging the values for different concentrations (right). **(D)** Summary of binding constants with the kinetic *k*_on_ determined for each experiment using K_D_ and *k*_off_.

**Figure S9:**
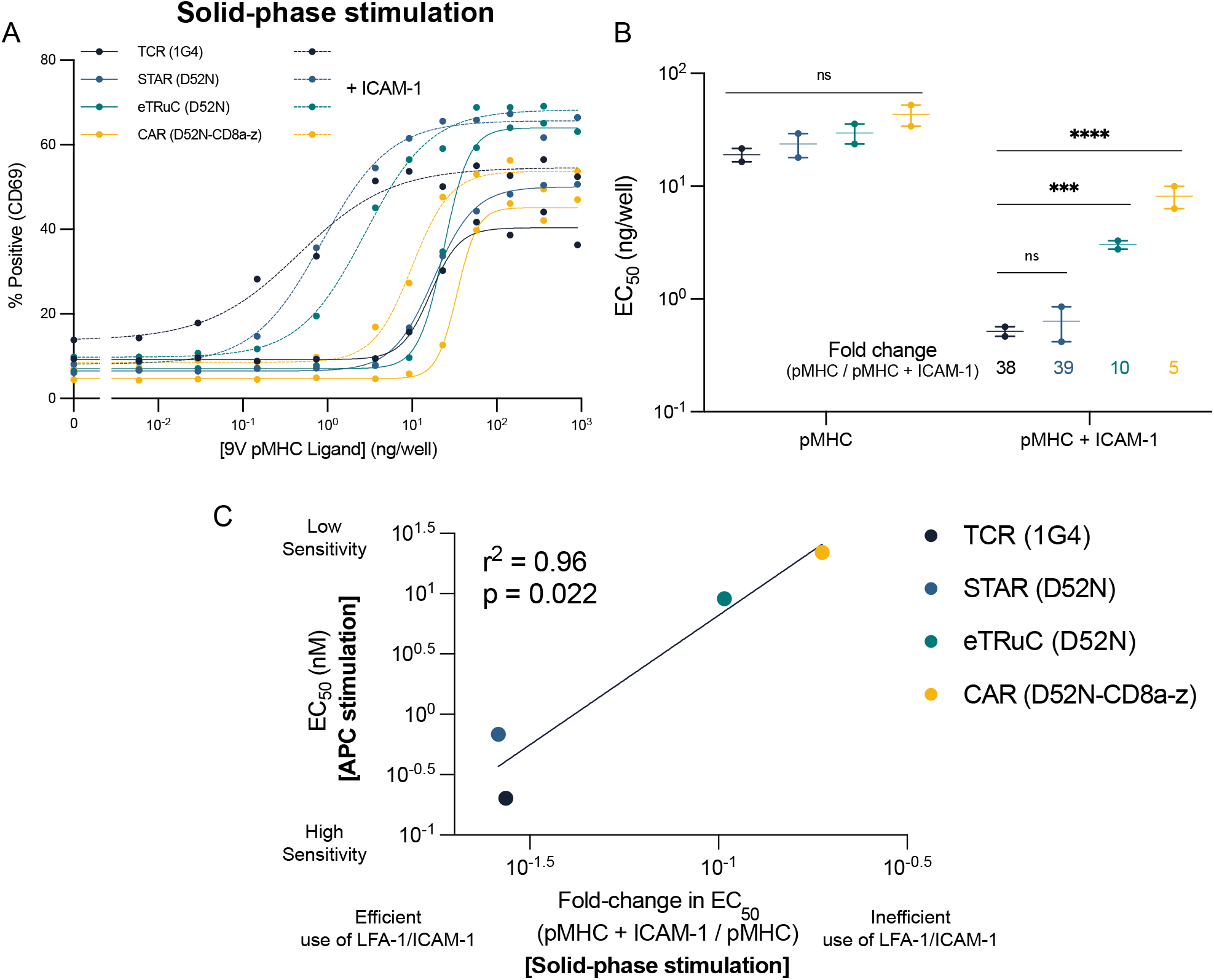
The ability of TCR-like chimeric antigen receptors to recapitulate the sensitivity of the TCR depends on the efficiency with which they are able to exploit the LFA-1 adhesion interaction. **(A)** T cells expressing the indicated antigen receptor were stimulated by a titration of purified pMHC alone (solid lines) or in combination with a fixed concentration of purified ICAM-1 (dashed lines). **(B)** Fitted EC_50_ values from two independent experiments. **(C)** The averaged EC_50_ values for CD69 upregulation from the APC stimulation assay (from Fig. 4C) are plotted over the averaged fold-change in EC_50_ for CD69 induced by the addition of ICAM-1 from the solid-phase stimulation assay (from panel B). The EC_50_ values are compared using a one-way ANOVA on log-transformed values (B). Abbreviations: *** = p-value≤0.001, **** = p-value≤0.0001.

